# Landscapes of transposable elements in *Apis* species by meta-analysis

**DOI:** 10.1101/2020.04.15.035063

**Authors:** Kakeru Yokoi, Kiyoshi Kimura, Hidemasa Bono

## Abstract

Transposable elements (TEs) are grouped into several classes with diverse sequences. Owing to their diversity, studies involving the detection, classification, and annotation of TEs are difficult tasks. Moreover, simple comparisons of TEs among different species with different methods can lead to misinterpretations. The genome data of several honey bee (*Apis*) species are available in public databases. Therefore, we conducted a meta-analysis of TEs, using 11 sets of genome data for *Apis* species, in order to establish the basal TE data (termed here as the ‘landscape of TEs’). Consensus TE sequences were constructed and their distributions in the *Apis* genomes were determined. Our results showed that TEs from several limited families mainly consisted of *Apis* TEs and that more DNA/TcMar-Mariner consensus sequences and copies were present in all *Apis* genomes tested. In addition, more consensus sequences and copy numbers of DNA/TcMar-Mariner were detected in *Apis mellifera* than in other *Apis* species. These results suggest that TcMar-Mariner might exert *A. mellifera*-specific effects on the host *A. mellifera* species. In conclusion, our unified approach enabled comparison of *Apis* genome sequences to determine the TE landscape, which provide novel evolutionary insights into *Apis* species.

**Simple Summary:** Studies on the detection of transposable elements and their annotations have posed several challenges. For example, simple comparisons of transposable elements in different species using different methods can lead to misinterpretations. Thus, assembling data for transposable elements analyzed by unified methods is important for comparison purposes. Therefore, we performed a meta-analysis of transposable elements identified using genome datasets from 5 *Apis* species (11 sets of genome data) and specific software to detect the transposable elements, which revealed the landscapes of transposable elements. We examined the types and locations of transposable elements in the *Apis* genomes. The landscapes of transposable elements showed that several limited transposable element families consisted mainly of *Apis*-associated transposable elements. These limited families include DNA/TcMar-Mariner and DNA/CMC-EnSpm. In addition, more DNA/TcMar-Mariner consensus sequences and copies were detected in *Apis mellifera* than in other *Apis* species. These data suggest that TcMar-Mariner might exert *A. mellifera*-specific effects in the host *A. mellifera* species. Our landscape data provide new insights into *Apis* transposable elements; furthermore, detailed analyses of our data could pave the way for new biological insights in this field.

## 1. Introduction

Transposable elements (TEs) are mobile DNA sequences that undergo change in their positions within a genome [1]. TEs occur in diverse forms and are found in the genomes of many species. Numerous effects of TEs on the host species have been reported. To mention some specific effects of TEs, they can serve as a source of mutations, lead to host-genome rearrangements, and change gene expression at the level of transcription. TEs can be divided into two classes: class I and class II (sometimes referred to as retrotransposons and DNA transposons, respectively) [1–3]. Class I TEs use an RNA intermediate and a “copy-and-paste” mechanism [1]. Class I TEs are further divided into subclasses (referred to as “order” in [3]), namely Long Terminal Repeats (LTRs), Dictyostelium intermediate repeat sequence (DIRS) and non-LTRs. LTRs are divided into several superfamilies (e.g. Copia, Gypsy and ERV) while non-LTRs are divided into other several superfamilies (e.g. Long Interspersed Nuclear Elements (LINEs), Short Interspersed Nuclear Elements (SINEs) and Penelope), several superfamilies of which some are divided into several families. Class II TEs move using a “cut-and-paste” mechanism through a DNA intermediate[1,3–5]; however, the Helitron type moves in a “peel-and-paste” manner [6]. Class II TEs are divided into subclasses (orders): Terminal inverted repeat (TIR) (possessing transposase in its coding region), Crypton, Helitron and Marverick. Each category is further classified into subfamilies, of which some are divided into several families [1,3]. For example, Tc1/Mariner is one of the subfamilies belonging to TIR subclass, and Tc1/Mariner is further classified into Tc1 or Mariner. The TE distributions and states of each species has specific features. Thus, performing a comparative analysis of the distributions and states of TEs among several species can potentially uncover some new insights into these species related to their TEs.

Honey bees, which belong to the Hymenoptera; Apidae, are important insects for honey production. They also pollinate wild plants and crops [7] and have been used as models of social insect species. Because of its widespread occurrence, whole genome sequencing of a representative honey bee species, the western honey bee (*Apis mellifera* [Am]), was completed at a very early phase among insect species [8]. This led to whole-genome sequencing of other honey bee species, including several Am and *A. cerana* sub-species. Genome data are currently available in public databases for the following honey bees: *A. cerana japonica* (Acj) [9], *A. cerana* Korean native (Ack) [10], *A. cerana* China native (Acc) [11], *A. dorsata* (Ad) [12], *A. florea* (Af), *A. laboriosa* (Al) [13], *A. mellifera carnica* (Carniolan honey bee) (Amcar), *A. mellifera intermissa* (Ami) [14], *A. mellifera caucasica* (Caucasian honey bee) (Amcau), and *A. mellifera* (German honey bee) (Amm) (Table 1). Am, *A. cerana*, Ad, Af are the 4 major *Apis* species [7]. Am and Acc genome data were recently updated using the long-read sequencer, and the N50 values have improved dramatically [15,16].

**Table 1.** *Apis* genome assemblies used in this study.

| Organism name [reference] | GenBank assembly accession ID | Genome size (bp) | Contig N50 | Abbreviation in this study |
| --- | --- | --- | --- | --- |
| *A. mellifera* [15] * | GCA_003254395.2 | 225,250,884 | 5,382,476 | Am |
| *A. cerana japonica* [9] | GCA_002217905.1 | 211,200,590 | 179,487 | Acj |
| *A. cerana Korea* native [10] | GCA_001442555.1 | 228,331,812 | 43,751 | Ack |
| *A. cerana China* native [16] * | GCA_011100585.1 | 215,670,033 | 3,898,192 | Acc |
| *A. dorsata* [12] | GCA_009792835.1 | 223,527,749 | 30,868 | Ad |
| *A. florea* | GCA_000184785.2 | 229,015,090 | 24,915 | Af |
| *A. laboriosa* [13] | GCF_014066325.1 | 226,078,798 | 303,790 | Al |
| *A. mellifera intermissa* [14] | GCA_000819425.1 | 243,566,977 | 504 | Ami |
| *A. mellifera* (German honey bee) * | GCA_003314205.1 | 227,036,473 | 5,131,172 | Amm |
| *A. mellifera carnica* (Carniolan honey bee) * | GCA_013841245.1 | 226,044,179 | 2,692,667 | Amcar |
| *A. mellifera caucasica* (Caucasian honey bee) | GCA_013841205.1 | 224,766,697 | 3,303,520 | Amcau |

Asterisks indicate chromosome-level genome assembly data according to NCBI genome assembly statistics in the NCBI dataset database (URL: https://www.ncbi.nlm.nih.gov/datasets/). See discussion section.

According to these *Apis* genome reports, *Apis* genomes contain relatively few TEs, which mainly consist of class II TEs, particularly Mariner-like-elements (MLEs), whereas some other representative insect genomes (e.g., silkworm *Bombyx mori* [17], yellow fever mosquito *Aedes aegypti* [18], and red flour beetle *Tribolium castaneum* [19,20]) contain higher numbers of TEs and MLEs.

Due to their ability to “transpose” within the genome, TEs have increased in number within the genome during evolution. In addition, new TEs enter the genome via horizontal transmission from other species. TE sequences have high diversity, due to the accumulation of mutations, which leads to many variants [1,21]. TE insertion and removal can indirectly cause rearrangements in host-genome sequences, leading to duplications or reshuffling around TEs in the host genome. These events can occur in genes or expression-regulation sites. Moreover, TEs can cause genome structural diversities long after TE could lose the capacity to move. Therefore, accurate TE detection and annotation are difficult to achieve. Although the basal TE status of each genome report available is important, comparisons among multiple species are suboptimal because the TE statuses were constructed using different methods and different software versions. Instead, new knowledge related to TEs could be obtained by studying the landscapes of TEs (the types of TEs and their positions in different *Apis* genomes), applying a unified TE analysis to the *Apis* genome data, and comparing the TE status between different species.

Comparing TE composition data among different species is important for genomic and evolutionary research, as indicated above. Recently, one report provided basal TE data for various insect species, suggesting that the content and diversity of TEs and genome sizes are related [22]. Other reports have provided evolutionary insights into *pogo* and *Tc1/mariner* by comparing the status data for these TE families in Apoidea genomes [23]. In this study, to obtain landscape data for such comparisons, a meta-analysis was performed using genome data from the 11 *Apis* genome data (5 *Apis* species) listed in Table 1, which are available in a public database (Figure 1). Specifically, we first performed *de novo* TE detection and then constructed consensus sequences for TEs with the same parameters, using RepeatModeler2 with the *Apis* genome data [24]. RepeatModeler2 runs multiple software packages to search for TEs and repetitive sequences, enabling accurate searches for TEs. To perform a detailed classification of the consensus sequences belonging to the Mariner or MLE family (the most prevalent among TE families in *Apis* genomes), phylogenetic analysis of MLE consensus sequences was performed. Finally, the distributions of repetitive elements, including the detected TEs, were investigated in all 11 *Apis* genomes using RepeatMasker, and the TE landscapes of *Apis* species were drawn. By comparing the TE statuses of different of *Apis* species and making use of the landscape data, we obtained new insights into TEs in *Apis* species

**Figure 1.**
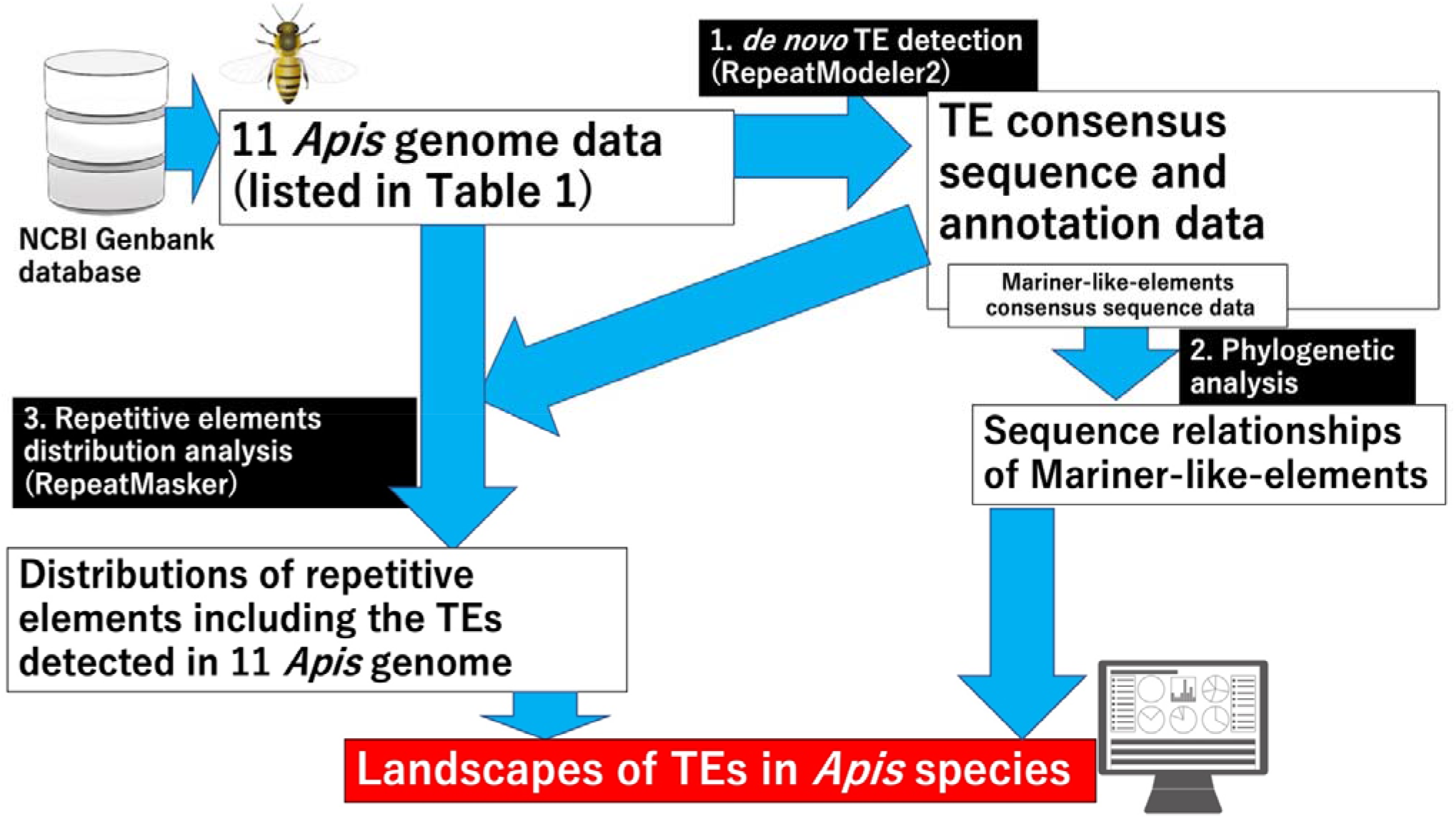
Workflow of the data analyses performed in this study. *De novo* TE detection was performed using 11 *Apis* genome sequences (Table 1) from NCBI genome database (URL: https://www.ncbi.nlm.nih.gov/genome/) using RepeatModeler2 [24]. Phylogenetic analysis revealed MLE relationships, where the most abundant consensus sequences were detected among the TE families in *Apis* species. The distributions of repetitive elements, including the TEs detected by RepeatModeler2, were investigated using RepeatMasker. The landscapes of TEs in *Apis* species were obtained using both sets of results, which led to new insights into TEs in *Apis* species. The images in Figure 1 were obtained from TogoTV (© 2016 DBCLS TogoTV).

## 2. Materials and Methods

### 2.1. Genome data used in this study

All genome data used in this study were downloaded from the NCBI Assembly section (https://www.ncbi.nlm.nih.gov/assembly/). The GenBank assembly accession IDs, genome sizes, N50 values, and abbreviations of each genome data point are presented in Table 1.

### 2.2. De novo detection of transposable elements consensus family sequences

*De novo* detection of TE consensus family sequences was performed using Repeat Modeler2 (version DEV) with the default settings and the genome data indicated in Table 1 [24].

### 2.3. Phylogenetic analysis

The detected TE sequences of some families were aligned using Clustal Omega (version 1.2.4) [25]. To construct approximately maximum-likelihood phylogenetic trees, aln files and Clustal Omega output files were further analyzed using FastTree (version 2.1.10) [26]. To visualize the phylogenetic trees, the FastTree output files (newick files) were loaded into MEGAX (version 10.1.7) [27].

### 2.4. Distribution analysis of repetitive elements in Apis genomes

The distributions of repetitive elements (including the TEs detected with RepeatModeler2) were investigated using the TE sequences as libraries and RepeatMasker (version 4.1.2-p1), with the default settings [28]

## 3. Results

### 3.1. Detection of transposable elements in Apis genomes

To determine the types of TEs in the *Apis* genomes, *de novo* TE detection was performed, and consensus TE family sequences were constructed with RepeatModeler2 using the *Apis* genomes shown in Table 1. The detection procedure used with RepeatModeler2 was described in detail previously [24]. Briefly, RepeatModeler2 runs different *de novo* repeat-detection programs such as RECON [29], RepeatScout [30], LtrHarvest [31] and Ltr_retriever [32]. The constructed family models from each software program are merged, redundancies are removed, and consensus sequences are constructed. The consensus sequences are annotated using RepeatClassifier, which compares the consensus sequences to several databases, including Dfam [24]. The output files from RepeatModeler2 are provided in Supplement data 1. The numbers of consensus sequences for each family are shown in Table 2. More consensus sequences were for class II TEs than for class I TEs. Among the class II TEs, DNA/TcMar-Mariner, that is MLE, DNA/TcMar-Tc1, DNA/hAT-Ac, DNA/CMC-EnSpm, and DNA/CMC-PiggyBac consensus sequences were constructed for all or 10 of the 11 *Apis* genomes studied, whereas the consensus sequences of other families were constructed in less than three *Apis* genomes. With class I TEs, the consensus sequences of three LTRs (LTR/Copia, LTR/Gypsy, and LTR/Pao) were constructed in more than 9 of the 11 *Apis* genomes.

**Table 2.**
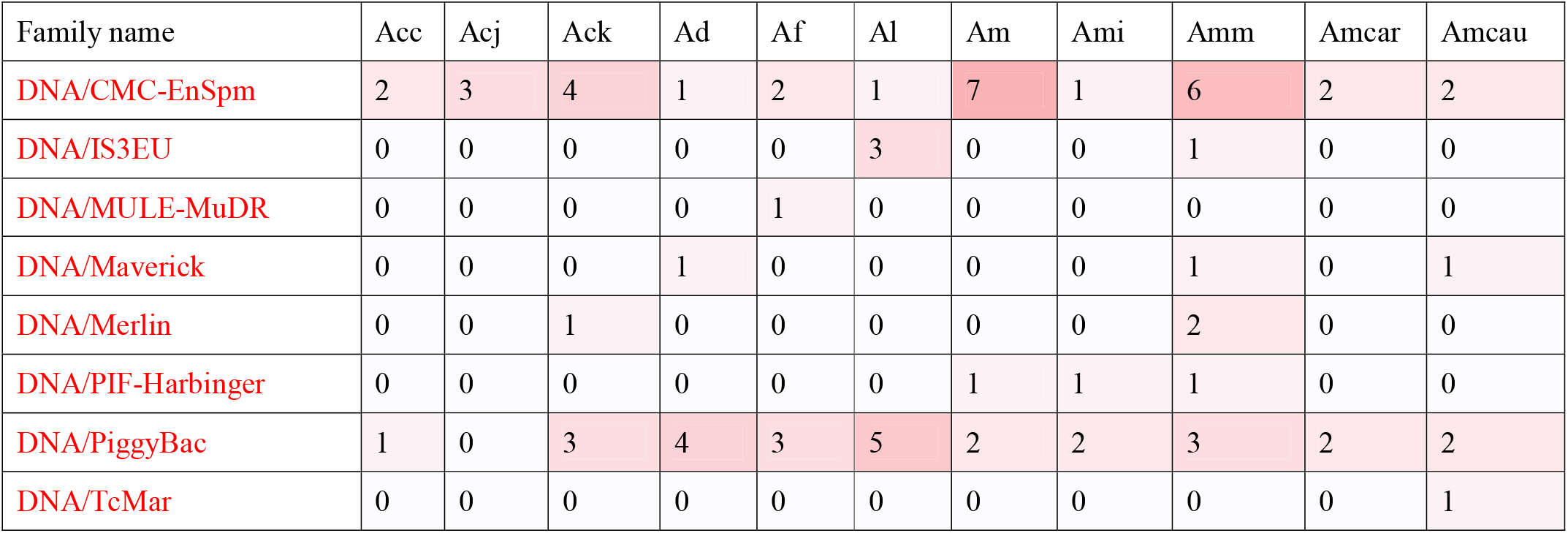
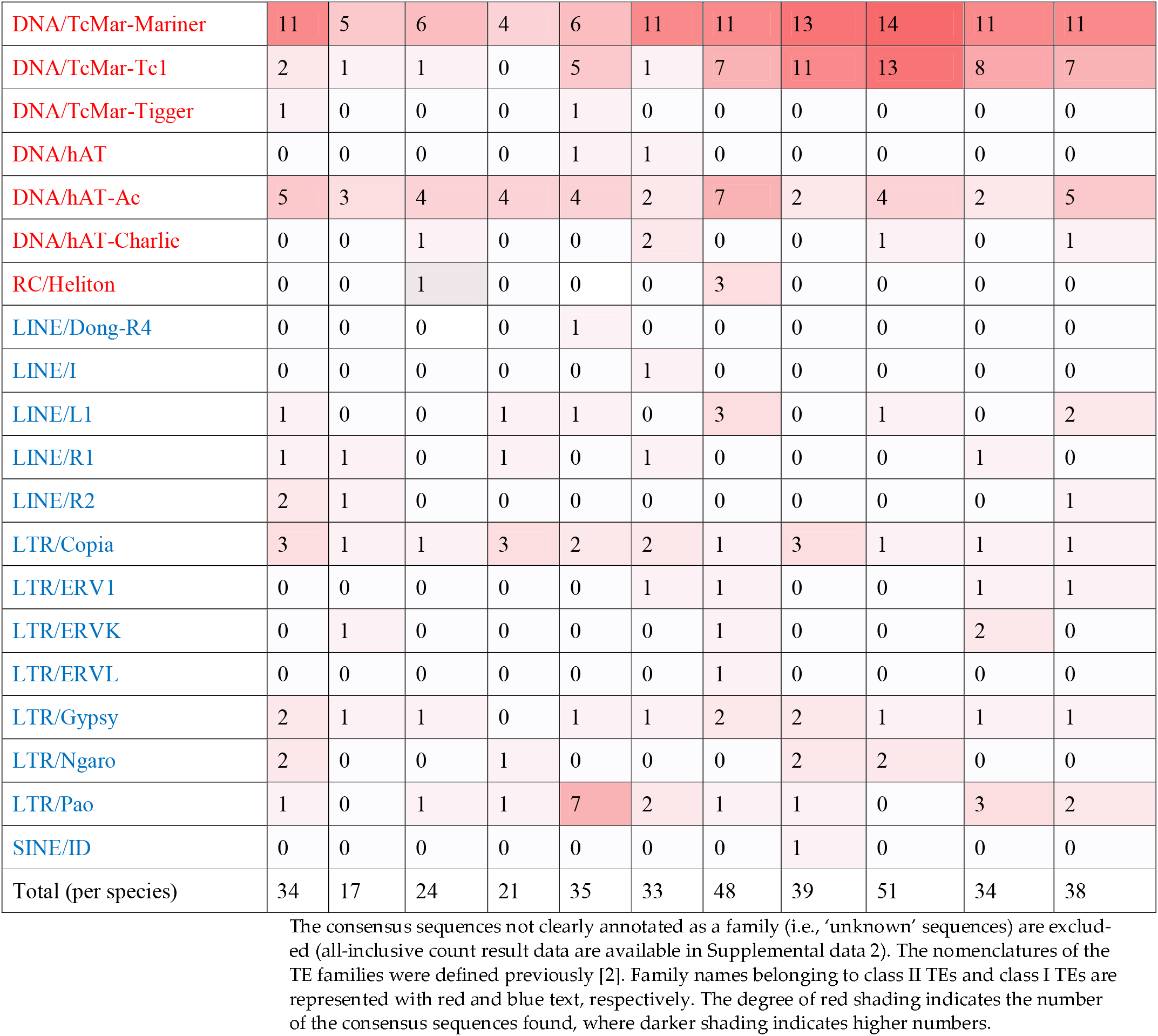
Total numbers of consensus sequences in the TE families of all *Apis* species, based on *de novo* TE detection with RepeatModeler2 [24].

Next, we investigated the differences in the numbers of consensus TE sequences among *Apis s*pecies. As shown in the “Total (per species)” row of Table 2, more consensus TE sequences were constructed with the *A. mellifera* species (Am, Ami, Amm, Amcar, and Amcau: 48, 39, 51, 34, and 38, respectively) than with the other Apis species (Acc, Acj, Ack, Ad, and Af: 34, 17, 24 21, and 35, respectively). Furthermore, more DNA/TcMar-Mariner sequences were detected with the *A. mellifera* species and Acc, representing the highest numbers of a consensus sequence constructed among the TE families. In addition, relatively high numbers of DNA/TcMar-Tc1 sequences were constructed with the *A. mellifera* species. These findings indicate that differences in the total number of consensus sequences among the *A. mellifera* species and other *Apis* species were mainly due to differences in DNA/TcMar-Mariner and DNA/TcMar-Tc1 consensus sequences.

### 3.2. Sequence analysis of TcMar-Mariner consensus sequences

As indicated in the previous section, the highest numbers of consensus sequences were constructed for the TcMar-Mariner family, among the TE families detected with RepeatModeler2. To obtain a more detailed classification, multiple sequence alignments were performed using the TcMar-Mariner consensus sequences (Supplemental data 3), Ammar1-6 (which were previously reported as *A. mellifera* MLEs [7]), and MLE consensus sequences of other species (mentioned in an MLE-related report [20]; Supplemental data 4), where the subfamilies have been annotated. Based on the alignment results, a phylogenetic tree was constructed (Figure 2 and Supplemental data 5 contain the raw data and related files). As shown in Figure 2 several clusters formed in the phylogenetic tree. MLEs annotated as a subfamily were expected to be located in a cluster; however, the phylogenetic tree showed that no MLEs belonging to a single subfamily were located in a single cluster. These results showed that the classifications of the MLE subfamilies, which are based on the amino acid sequences of transposase in MLEs, conflicted with the results of the nucleotide sequence-based analyses we performed. All-inclusive count result data are available in Supplemental data 2

**Figure 2.**
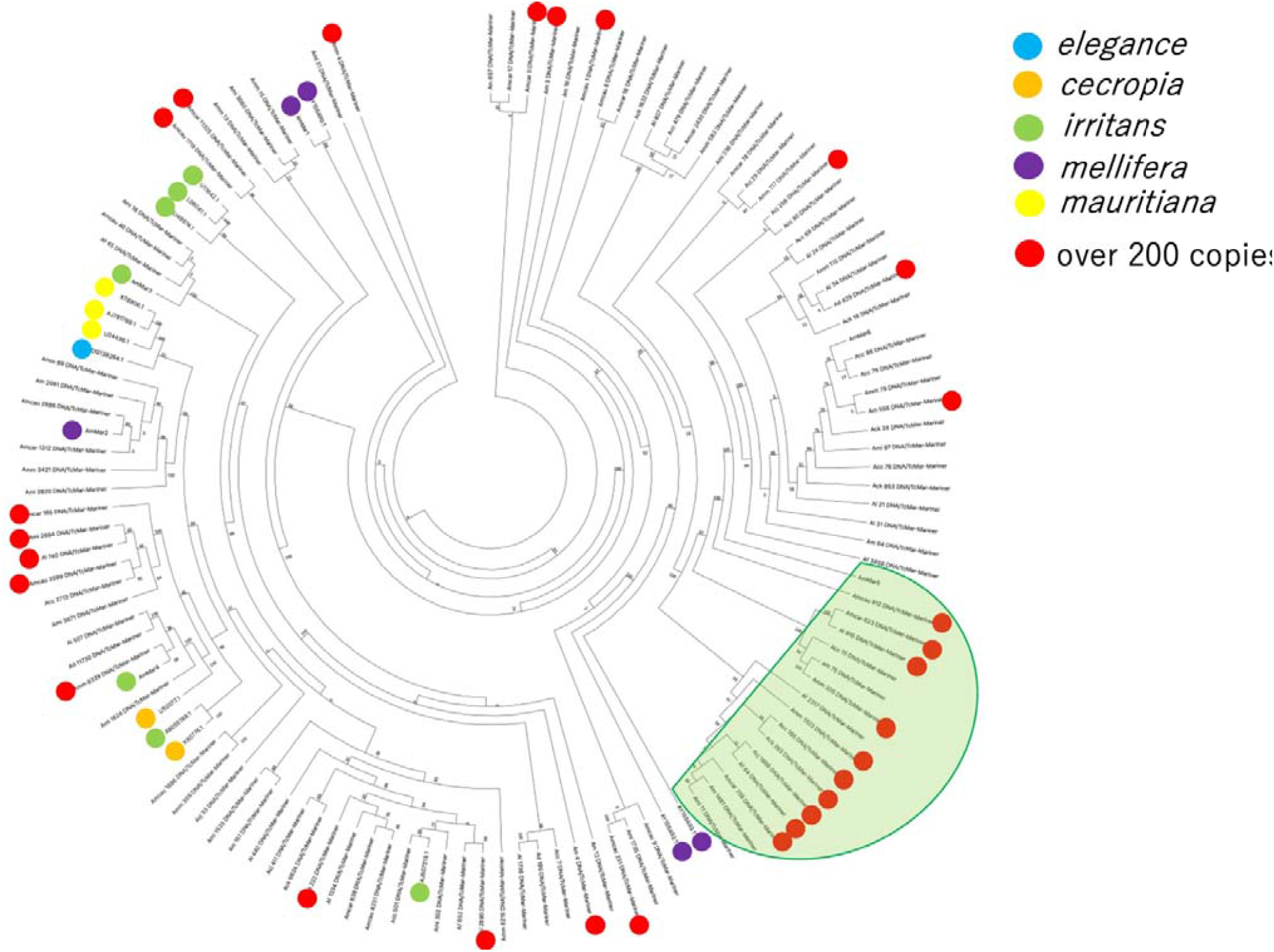
Phylogenetic tree of *Apis* TcMar–Mariner consensus sequences identified in this study. The MLE sequences of other species and *A. mellifera* were annotated with Mariner subfamilies in previous reports [8,20]. Blue, orange, green, purple, and yellow circles located at end of each node (MLE sequences from the previous reports) indicate the MLE subfamilies. The red circles indicate consensus sequences detected with more than 200 copies. The green semicircular shading encompasses a clade including many sequences with over 200 copies. The numbers at the branches indicate bootstrap values. A high-resolution phylogenetic tree data is available in Supplemental data 5.

### 3.3. Distribution analysis of transposable elements in Apis genome

To determine the distributions of the TEs detected with RepeatModeler2 in *Apis* genomes (Section 3.1), we ran RepeatMasker with the *Apis* reference genome as the input data (Table 1) and the consensus TE sequence data as libraries using Repeat Modeler2 (Supplemental data 1). RepeatMasker was used to screen the TE sequences (registered in Dfam or Repbase) or the consensus sequences as input data (mainly from RepeatModeler2) and simple repeat sequences as genomic query data (for greater detail, see [28]). Because of these software features, high numbers of short TE sequences were detected. The output files are provided in Supplemental data 6. The percentages of repetitive elements, including TEs, present in the *Apis* genomes are shown in Table 3. Our findings indicate that repetitive elements comprised approximately 7 to 12% of the *Apis* genome regions. The *A. mellifera* genomes (except for Ami) had higher percentages than the other *Apis* genomes. The percentages in the *Apis* genomes were lower than those reported for other insect species. (e.g., approximately 46.8% for *B. mori* [17], 20% for *T. castaneum* [20], 65% for *A. aegypti* [16], and 20% for *D. melanogaster* [20]).

**Table 3.**
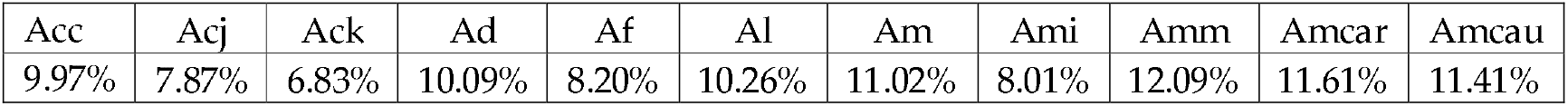
Percentages of repetitive elements present in each *Apis* genome.

The RepeatMasker results are summarized in .tbl files (RepeatMasker output files) and are available in Supplemental data 6. Because the summary files did not show the number of copies in the individual TE families, we counted them using .out files and other RepeatMasker output files (Supplemental data 6). The number of copies belonging to the TE families that were clearly annotated as a TE family member (e.g., DNA and SINE?) plus other repetitive elements (e.g., Simple repeat) in all *Apis* genomes are given in Supplemental data 7. The total copy numbers of class II and class I TE families in all *Apis* genomes are shown in Table 4 and Table 5, respectively. Overall, several TE families had multiple copies. Among these TE families, the TEs of class II had many more copies than those of class I. With regard to class II, more total number of copies (except for Ami) were observed in *A. mellifera* genomes than in the other *Apis* genomes (Table 4). In contrast, among the class I TEs, Acj, Ack, and Ami showed lower copy numbers, whereas Am had a higher copy number than the other *Apis* species (Table 5).

**Table 4.** Total copy numbers of class II TE families in the *Apis* genomes listed Table 1.

| Family name | Acc | Acj | Ack | Ad | Af | Al | Am | Ami | Amm | Amcar | Amcau |
| --- | --- | --- | --- | --- | --- | --- | --- | --- | --- | --- | --- |
| DNA/CMC-EnSpm | 1387 | 1684 | 1797 | 880 | 1060 | 692 | 2761 | 538 | 2200 | 1305 | 1518 |
| DNA/IS3EU | 0 | 0 | 0 | 0 | 0 | 107 | 0 | 0 | 169 | 0 | 0 |
| DNA/MULE-MuDR | 0 | 0 | 0 | 0 | 477 | 0 | 0 | 0 | 0 | 0 | 0 |
| DNA/Maverick | 0 | 0 | 0 | 165 | 0 | 0 | 0 | 0 | 59 | 0 | 193 |
| DNA/Merlin | 0 | 0 | 107 | 0 | 0 | 0 | 0 | 0 | 335 | 0 | 0 |
| DNA/PIF-Harbinger | 0 | 0 | 0 | 0 | 0 | 0 | 406 | 59 | 698 | 0 | 0 |
| DNA/PiggyBac | 138 | 0 | 316 | 845 | 474 | 826 | 456 | 318 | 848 | 797 | 678 |
| DNA/TcMar | 0 | 0 | 0 | 0 | 0 | 0 | 0 | 0 | 0 | 0 | 364 |
| DNA/TcMar-Mariner | 1254 | 630 | 798 | 631 | 903 | 1343 | 1892 | 1495 | 2641 | 2478 | 3475 |
| DNA/TcMar-Tc1 | 618 | 159 | 110 | 0 | 313 | 608 | 1010 | 1507 | 1656 | 1461 | 2300 |
| DNA/TcMar-Tigger | 230 | 0 | 0 | 0 | 118 | 0 | 0 | 0 | 0 | 0 | 0 |
| DNA/hAT | 0 | 0 | 0 | 0 | 98 | 201 | 0 | 0 | 0 | 0 | 0 |
| DNA/hAT-Ac | 657 | 510 | 233 | 821 | 673 | 409 | 1702 | 404 | 974 | 351 | 1736 |
| DNA/hAT-Charlie | 0 | 0 | 188 | 0 | 0 | 642 | 0 | 0 | 466 | 0 | 447 |
| RC/Heliton | 0 | 0 | 38 | 0 | 0 | 0 | 2852 | 0 | 0 | 0 | 0 |
| Total (per species) | 4284 | 2983 | 3587 | 3342 | 4116 | 4828 | 11079 | 4321 | 10046 | 6392 | 10711 |

The total numbers of TE families (detected using RepeatModeler2) were calculated using output files from RepeatMasker. The degree of red shading reflects the copy numbers found, where darker shading indicates higher copy numbers.

**Table 5.** Total copy numbers of class I TE families in the *Apis* genomes listed in Table 1.

| Family name | Acc | Acj | Ack | Ad | Af | Al | Am | Ami | Amm | Amcar | Amcau |
| --- | --- | --- | --- | --- | --- | --- | --- | --- | --- | --- | --- |
| LINE/Dong-R4 | 0 | 0 | 0 | 0 | 100 | 0 | 0 | 0 | 0 | 0 | 0 |
| LINE/I | 0 | 0 | 0 | 0 | 0 | 24 | 0 | 0 | 0 | 0 | 0 |
| LINE/L1 | 26 | 0 | 0 | 121 | 341 | 0 | 654 | 0 | 480 | 0 | 261 |
| LINE/R1 | 74 | 57 | 0 | 81 | 0 | 161 | 0 | 0 | 0 | 75 | 0 |
| LINE/R2 | 332 | 51 | 0 | 0 | 0 | 0 | 0 | 0 | 0 | 0 | 249 |
| LTR/Copia | 829 | 82 | 101 | 749 | 354 | 466 | 257 | 321 | 318 | 109 | 268 |
| LTR/ERV1 | 0 | 0 | 0 | 0 | 0 | 217 | 419 | 0 | 0 | 350 | 75 |
| LTR/ERVK | 0 | 356 | 0 | 0 | 0 | 0 | 326 | 0 | 0 | 1316 | 0 |
| LTR/ERVL | 0 | 0 | 0 | 0 | 0 | 0 | 52 | 0 | 0 | 0 | 0 |
| LTR/Gypsy | 574 | 46 | 44 | 0 | 147 | 233 | 1000 | 426 | 417 | 203 | 499 |
| LTR/Ngaro | 153 | 0 | 0 | 91 | 0 | 0 | 0 | 48 | 483 | 0 | 0 |
| LTR/Pao | 44 | 0 | 228 | 416 | 730 | 213 | 677 | 57 | 0 | 300 | 1237 |
| SINE/ID | 0 | 0 | 0 | 0 | 0 | 0 | 0 | 24 | 0 | 0 | 0 |
| Total (per species) | 2032 | 592 | 373 | 1458 | 1672 | 1314 | 3385 | 876 | 1698 | 2353 | 2589 |

The total numbers of TE families (detected using RepeatModeler2) were calculated using output files from RepeatMasker. The degree of red shading indicates the number of the consensus sequences found, where darker shading indicates higher numbers.

Among the class II TE families, copies of DNA/CMC-EnSpm, DNA/TcMar-Mariner, and DNA/hAT-Ac were detected in all *Apis* genomes tested (Table 4). Over 1000 copies of DNA/TcMar-Mariner were detected in all *A. mellifera* species and in Acc and Al. In addition, 1000 DNA/CMC-EnSpm copies were detected in all *Apis* species tested, except for Ad, Al and Ami, whereas 1000 copies of DNA/hAT-Ac were detected in Am and Amcau genomes. In the case of DNA/TcMar-Tc1, over 1000 copies were detected in all *A. mellifera* species, but no copies were detected in Ad. Over 400 DNA/PiggyBac copies were detected in some *Apis* genomes, but no copies were detected in the Acj genome. Among the class I TE families, copies of LTR-Copia were detected in all *Apis* genomes tested, and copies of LTR-Gypsy and LTR/Pao were detected in 10 and 9 *Apis* species, respectively.

As shown above, abundant copies of DNA/TcMar-Mariner and DNA/CMC-EnSpm were detected in all *Apis* genomes tested. To investigate this phenomenon in greater detail, the copy numbers of both TE families in each of the *Apis* genomes are shown in graphically in Figure 3. In the case of DNA/TcMar-Mariner, *A. mellifera* species (especially Amm, Amcar, and Amcau) had higher copy numbers than other *Apis* species (Figure 3A). In the case of DNA/CMC-EnSpm, Am and Amm had higher copy numbers than other *Apis* species, whereas Ad, Af, Al and Ami had fewer copies (Figure 3B). We further investigated which consensus TcMar-Mariner sequences in particular had many copies (Table 2). As shown in Figure 2, consensus sequences with more than 200 copies (indicated with red circles) were scattered over the trees, and several sequences with red circles were located in a single clade (represented with the green semicircular object). This clade contained the sequences of all *Apis* species tested, except for Af.

**Figure 3.**
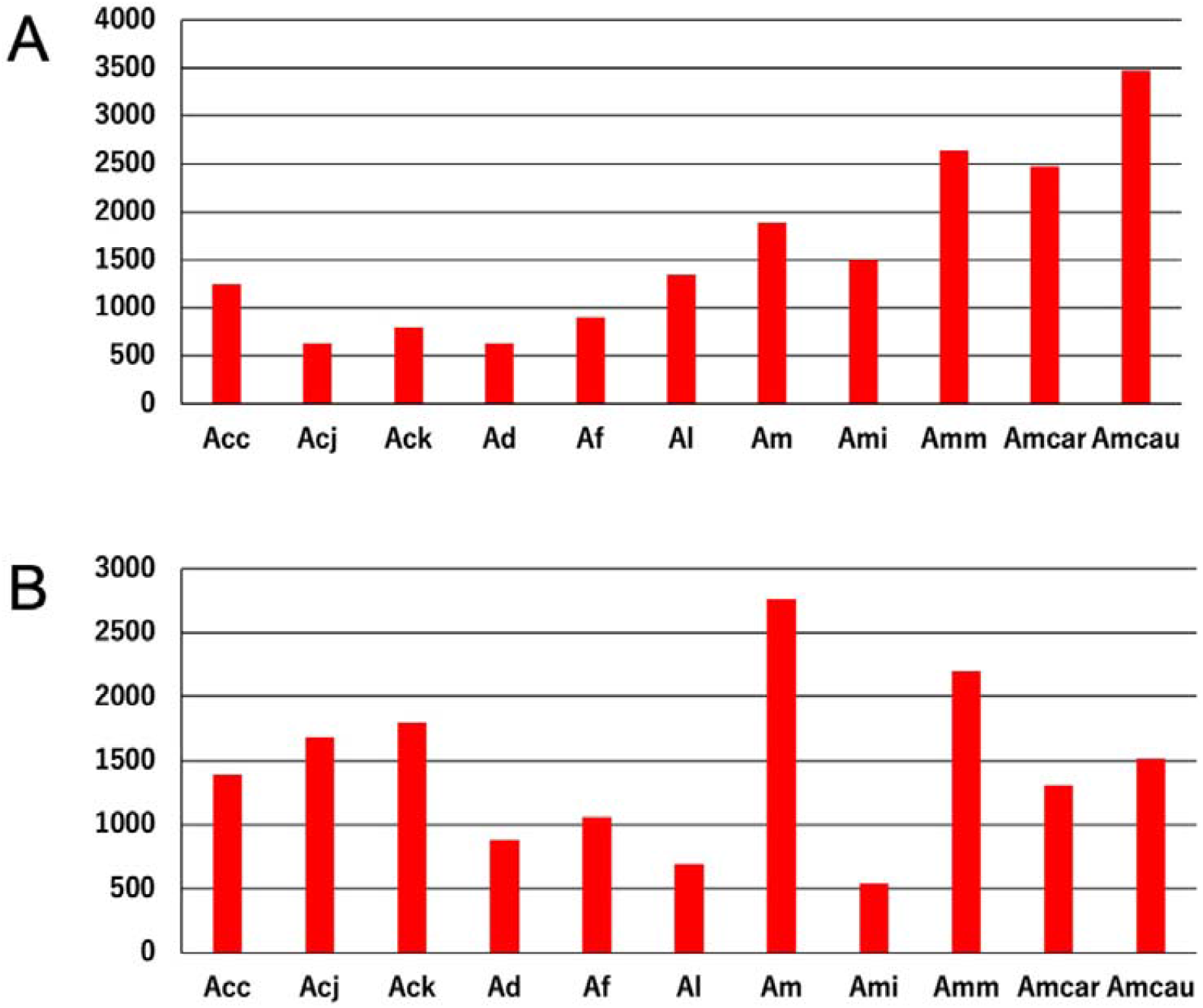
The total numbers of DNA/TcMar-Mariner (A) and DNA/CMC-EnSpm (B) TEs in each *Apis* genome listed in Table 1. Both TE families were detected using Repeat Modeler 2 and the total numbers of TEs were calculated using .out files from Repeat Masker. Abbreviations of names of *Apis* species in the figure are shown in Table 1.

## 4. Discussion

In this study, we investigated the landscapes of TEs in *Apis* species using *Apis* genome data, which are available in public databases, and TE consensus sequences were constructed. Sequence analysis was performed, and phylogenetic trees were constructed to reveal more detailed relationships for the MLEs, the consensus sequences of which are the most diverse among the TE families detected. Consequently, the distributions of repetitive elements, including the constructed consensus TE sequences within the corresponding *Apis* genomes, were revealed. Our landscapes showed that several limited TE families are mainly found in *Apis* genomes.

As described above, detecting TEs in genome sequences is a difficult task because TE sequences have many variants and deletions [21]. Therefore, the results related to TEs can be varied can vary when different methods are adopted. Our meta-analysis was performed using two major software packages that are commonly employed. RepeatModeler2 is commonly used for *de novo* TE detection with genome data [24]. This software package can also be used to construct consensus sequences. Multiple repeat-searching programs can be run, and merging the results of the program enables accurate detection of TEs (for benchmarking the results, see a previous article that described RepeatModeler2 [24]). RepeatMasker searches for simple repeats and TEs in queried genome data, with consensus sequences serving as the input data [28]. A series of analyses can provide accurate and comprehensive landscape data for TEs in the queried genome. These landscape data can be utilized for further detailed analyses (e.g., comparing the TE status between different species).

The genome assembly level of genome data can affect TE detections. As shown in Table 2, over half of *Apis* genome data we used from the public database are not “chromosome-level genome assembly data”. TE data in detailed points using scaffold-level genome assembly data and chromosome-level genome assembly data in the same species can be different. However, we think that the genome assembly level can not affect on main features of *Apis* TEs we showed; for example, several limited transposable element families consisted mainly of TEs and much more copies of DNA/CMC-EnSpm, DNA/TcMar-Mariner in all *Apis* species genomes, which included various genome assembly levels. It is very interesting to analyze the two levels separately. However, since our goal is to clarify the landscape of TEs in the genus *Apis*, which is not affected by differences in genome assembly, we did not analyze them separately in this study.

There are adequate and reliable software for *de novo* TE detection in genome sequences, such as EDTA [33] and REPET [34]). The benchmarking results showed that RepeatModeler2 produced the output file which were similar to the curated libraries using several model species genome data, and showed better status related to the detected family quality and the detected sequences of fragmentation and redundancy than other software tested while these software showed better status related to some cases [24]. Considering these result, we decided to choose RepeatModeler2 for *de novo* TE detection.

As mentioned in the introduction part, there are more than 10 known species of honey bees, and by examining the four main species (Af, Ad, *A. cerana* and Am) and one closely related species (Al) [7] with RepeatModeler2 and RepeatMasker, which were used for *de novo* TE detections and revealing distributions of the detected TE families respectively, we were able to characterize the TEs common to the genus *Apis* without using all species, thus providing a “landscape” of the TEs in the genus *Apis*, which is our goal of this article. Interestingly, although Ad and Al are closely related species, the landscapes showed that there were several different features of TEs between the two species.

With both class II and class I TEs, several limited families have diverse consensus sequences, whereas the other families had a few consensus sequences in *Apis* genomes, implying that these limited TE families might exert several effects on host *Apis* species through several mechanisms (e.g., gene insertions or alterations at the transcription level) [35,36]. Comparisons of the consensus TE sequences among *Apis* species revealed that more consensus sequences were constructed for *A. mellifera* than for the other *Apis* species, which was mainly due to DNA/Tc-Mariner and DNA/Tc-Tc1 (which have many consensus sequences). These results suggest that some of the TEs could have had effects on *A. mellifera* species that might not have occurred in other *Apis* species.

Among the several characteristics of honey bee TEs revealed by the landscape data, it is worth noting the patchy distribution of each TE. Some TEs are identified only in certain *Apis* species. For example, DNA/MULE-MuDR was only found only in Af and TcMar-Tigger was found only in Acc. Also, RC/Heliton was found only in Am and not is any other Am subspecies. This biased and patchy distribution of the TEs is well known in other species [22,23]. The most famous example of such a distribution is the P element, which is present only in certain strains (e.g. P strains) of *Drosophila melanogaster* [37]. Using this landscape data, we plan to conduct such detailed comparative analysis in the future.

Many Mariner or MLE consensus sequences were constructed for *Apis* species in this study. As described above, these consensus sequences were constructed using RepeatModeler2, which runs repeat detection programs and annotates the constructed sequences using several databases including Dfam [2,24]. A further detailed classification of these MLEs was performed. This was done by generating alignments and constructing phylogenetic trees using MLE consensus nucleotide sequences that were previously annotated with MLE consensus sequences. Our results revealed that MLE sequences annotated as part of the same MLE subfamily did not form a single clade. MLEs, which have a DD34D catalytic motif in their encoded transposase, are classified into subfamilies based on their transposase amino acid sequences [38,39]. This classification principle must be respected; however, we believe that nucleotide-based classification may also be required. As shown in this study, an enormous number of TE nucleotide sequences can be detected in target genomes because the whole-genome data of many species are available in public databases, and sophisticated TE detection software, such as RepeatModeler2, are now available. [24]. Some detected MLEs do not encode transposases of sufficient length because of mutations or deletions in their sequences. Therefore, annotations based on amino acid sequences cannot be used to study such MLE sequences. According to a previous report, annotation methods for studying subfamilies are fraught with problems such as a lack of reproducibility [21]. The development of nucleotide-based annotations of MLE subfamilies is essential for future genome analysis, and our data could lead to future research in this field.

Nucleic acid-based analysis using RepeatModeler2 and RepeatMasker (in this study), and analysis using the consensus amino acid sequence of transposase have yielded several different results [22,23]. However, even if the same genome data are used for nucleic acid-based analysis, the results will differ slightly depending on the method used, and the number of each TE found differs depending on the software used. For example, as mentioned above, we identified many copies of many DNA/TcMar-Tc1 types. However, in a previous analysis using the tblastn method with amino acid sequences against Apoidea genomes, including some *Apis* genomes [23], these TEs were not found in the *Apis* genomes. This discrepancy may reflect our method used, which recognizes the Tc1 and Mariner types as different, whereas the previous analysis considered them to be the same type of TEs. Another example is the detection of DNA/CMC-EnSpm in all *Apis* genomes tested, whereas previous findings indicated that DNA/CMC-EnSpm was absent from the Am genome [40]. This may be because the TEs annotated as DNA/CMC-EnSpm in this study were classified as putative elements, unclassified, or classified Class II TEs. Indeed, *Nasonia vitripennis* DNA/CMC-EnSpm was registered in Repbase (e.g., EnSpm-2_NVi) [41], and another report showed that CMC TEs were detected in the Am genome [22]. This discrepancy illustrates the difficulty of classifying TEs. However, our landscape was successful in providing a general framework for the TEs of the *Apis* genus. Further evolutionary studies of TEs will require analysis of the individual TEs found. Recent advances in bioinformatics have made this possible.

RepeatMasker results for the *Apis* species showed that repetitive elements comprise approximately 7 to 12% of *Apis* genomes, which is lower than that of many other insect species [20]. However, these percentages are consistent with previous reports [9,10,12,14–16], which validate the accuracy of our datasets and the analytical methods used in this study. Comparing the numbers of TEs among *Apis* species showed that *A. mellifera* species, with the exception of Ami, have more TEs than other *Apis* species. Ami showed lower percentages of repetitive elements, perhaps because the N50 value of Ami was much lower than those of other *Apis* species. Thus, we conclude that *A. mellifera* species have more repeat regions and TEs than other *Apis* species.

The total number of TE copies in each TE family showed that families with a higher number of consensus sequences had a higher number of elements. In addition, more copies of class II TEs than class I TEs were detected. Furthermore, these results revealed that the TEs of several limited families in both classes (II and I) consisted of *Apis* TEs. Most of these results have the same tendencies as those of the consensus TE sequences. These results suggest that TEs belonging to limited TE families mostly consist of *Apis* TEs. A more detailed investigation also revealed more class II TEs in *A. mellifera* genomes, except for Ami, than in other *Apis* genomes. TcMar-Mariner/MLEs were identified as a family with a high number of copies in all *Apis* species tested. The phylogenetic tree revealed that, although several MLE consensus sequences of all *Apis* species tested (except for Af, which had over 200 copies) were located in a clade, these sequences were scattered in the trees, suggesting that the abundant MLEs may have been copied from many consensus sequences rather than from a very limited number of consensus sequences.

Although clear differences were found in the number and type of TEs between species, it is interesting to note that variation has occurred within species. This may be due to differences in the quality of the genome data. Among the genome data used in this study, the Ami genome data showed a much lower contig N50 number than the other genome data. No significant correlations were found between the contig N50 numbers for the *Apis* genome data and the numbers and types of TEs. It would be interesting from an evolutionary point of view if the intraspecific variation observed here was not due to differences in the quality of the genome data. Our findings indicate that many TEs increase in number, shift, or propagate horizontally in the genome after subspeciation. Further studies are required to elucidate these differences.

In this study, we performed a meta-analysis of *Apis* TEs using *Apis* whole-genome data and TE-detection software. Through this analysis, we determined basal data for TEs showing the specific types of TEs and their positions in the *Apis* genomes (the *Apis* TE landscapes). We also showed that several limited TE families exist in *Apis* genomes and that *A. mellifera* species have more TEs, mainly due to MLEs. The findings of this study provide several new insights into the genomes of *Apis* species. The landscape data obtained in this study can be compared to TE data for other species, including Hymenoptera or other insects [20,22,23], leading to findings related to the evolution of TEs between these species. In addition, analyzing our landscape data in greater detail could help elucidate new TE-related biological insights for *Apis* species.

## Supplementary Materials

All supplemental data are available in figshare (DOI: 10.6084/m9.figshare.c.5847335).

**Supplemental data 1** Output files (fasta file and stk file) of RepeatModeler2. Out files of family consensus files of transposable elements by RepeatMolder2. {abbreviation of *Apis* species}.fa and {abbreviation of *Apis* species}.stk contain consensus sequences with metadata describing transposable element families. Nomenclature of them is shown in [2] (DOI: 10.6084/m9.figshare.19189004).

**Supplemental data 2** Numbers of transposable element consensus sequences of all families constructed in all the *Apis* genomes tested by RepeatModeler2 (DOI: 10.6084/m9.figshare.19189127).

**Supplemental data 3** Consensus sequences annotated as Mariner in *Apis* genomes by RepeatModeler2. All sequences were extracted from RepeatModeler2 output fasta files. Abbreviations of *Apis* species are shown in table 1 (DOI: 10.6084/m9.figshare.19189055).

**Supplemental data 4** Consensus Mariner sequence files from other papers used for phylogenetic tree analysis. Ammar1-6 (Ammar.fa) are listed in Supplemental information of [8]. Other sequences (MLE.otherspecies.fa) are shown in Additional file 1 of [20] and Genbank ID of each sequence is shown in a description part (DOI: 10.6084/m9.figshare.19189073).

**Supplemental data 5** Phylogenetic analysis-related data. MLE_tree_ana.fa is Input data including all MLE consensus sequences plus Ammar and other species shown in (all_Mariner.fa, Ammar.fa and MLE.otherspecies.fa). MLE_tree_ana.aln is Clustal omega output file (aln), and MLE_tree_ana.newick is FastTree output data (newick). Trees.png is a high-resolution phylogenetic tree picture (DOI: 10.6084/m9.figshare.19189181).

**Supplemental data 6** Output files of RepeatMasker using TE consensus sequence files by RepeatModeler2 and *Apis* genome sequences. (DOI: 10.6084/m9.figshare.19189292).

**Supplemental data 7** Copy numbers of TEs in *Apis* genomes. Numbers of TEs which are counted by using RepeatMasker out files in Supplemental data 6 (DOI: 10.6084/m9.figshare.19189376).

## Author Contributions

Conceptualization, K.Y., K.K. and H.B.; methodology, K.Y., K.K. and H.B.; data validation, K.Y.; formal data analysis, K.Y.; data curation, K.Y., K.K. and H.B.; writing— original draft preparation, K.Y.; writing—review and editing, K.Y., K.K, and H.B.; supervision, K.Y.; project administration, K.Y.; funding acquisition, K.Y and H.B. All authors discussed the data and have read and agreed to the published version of the manuscript.

## Institutional Review Board Statement

Not applicable.

## Data Availability Statement

All data in this study are available in figshare as described in “Supplementary Materials”.

## Acknowledgments

We express our gratitude to Drs. Masatsugu Hatakeyama and Yudai Masuoka for their valuable discussions.

